# The protein translation machinery is expressed for maximal efficiency in *Escherichia coli*

**DOI:** 10.1101/802470

**Authors:** Xiao-Pan Hu, Hugo Dourado, Martin J. Lercher

## Abstract

Protein synthesis is the most expensive process in fast-growing bacteria^1,2^. The economic aspects of protein synthesis at the cellular level have been investigated by estimating ribosome activity^3–5^ and the expression of ribosomes^3,6^, tRNA^7–9^, mRNA^2^, and elongation factors^10,11^. The observed growth-rate dependencies form the basis of powerful phenomenological bacterial growth laws^5,12–16^; however, a quantitative theory allowing us to understand these phenomena on the basis of fundamental biophysical and biochemical principles is currently lacking. Here, we show that the observed growth-rate dependence of the concentrations of ribosomes, tRNAs, mRNA, and elongation factors in *Escherichia coli* can be predicted accurately by minimizing cellular costs in a detailed mathematical model of protein translation; the mechanistic model is only constrained by the physicochemical properties of the molecules and requires no parameter fitting. We approximate the costs of molecule species through their masses, justified by the observation that cellular dry mass per volume is roughly constant across growth rates^17^ and hence represents a limited resource. Our results also account quantitatively for observed RNA/protein ratios and ribosome activities in *E. coli* across diverse growth conditions, including antibiotic stresses. Our prediction of active and free ribosome abundance facilitates an estimate of the deactivated ribosome reserve^14,18,19^, which reaches almost 50% at the lowest growth rates. We conclude that the growth rate dependent composition of *E coli*’s protein synthesis machinery is a consequence of natural selection for minimal total cost under physicochemical constraints, a paradigm that might generally be applied to the analysis of resource allocation in complex biological systems.

## Introduction

Protein translation is central to the self-replication of biological cells. It is the energetically most expensive process in fast growing *E. coli* cells, accounting for up to 50% of the proteome^2^ and 2/3 of cellular ATP consumption^1^. It is likely that natural selection acted to optimize the efficiency of this central process. But what exactly is “efficiency” in the evolutionary context? In the late 1950s, it was hypothesized that ribosomes operate at a constant, maximal rate^3,4^, consistent with the observed linear dependence of ribosome concentration on growth rate^3,12,20,21^. This hypothesis was later proven untenable, as the activity of ribosomes was observed to increase with growth rate^8^. Klumpp *et al*.^5^ suggested that optimal translational efficiency corresponds to the parsimonious usage of translation-associated proteins, most notably ribosomal proteins, elongation factor Tu, and tRNA synthetases. While these authors were able to fit a coarse-grained phenomenological model to the data, their suggested evolutionary objective could also not explain the observed growth rate dependencies quantitatively (see **Supplementary Notes 1** for a discussion of Ref. ^5^). Thus, it is currently unclear to what extent translation has indeed been optimized by natural selection, and – if such optimization indeed occurred – whether its action can be expressed in terms of a simple objective function.

Here, we propose an entirely different evolutionary objective, based on the experimental observation that cellular dry mass per cell volume is approximately constant across environments and growth rates in *E. coli*^17^, as is the total mass concentration in the cytosol^22^. If the cell allocates more of this limited mass concentration “budget” to one particular process, less is available to other processes. The upper bound for the cytosolic mass concentration, beyond which diffusion becomes inefficient, is a fundamental constraint on cellular growth^23,24^, and we thus use the cytosolic mass concentration of a particular molecule type as an approximation to its cost.

We hypothesize that to maximize the *E. coli* growth rate in a given environment, natural selection minimizes the total cost of translation components utilized to achieve the required protein production rate. An analogous optimality principle has been used to understand the relationship between enzyme and substrate concentrations, explaining the scaling of *E. coli* proteome sectors with growth rate^34^. We emphasize that the optimal efficiency of the translation machinery is not based on the maximization of ribosome activity, but on the minimization of the combined cost of the complete translation machinery at a given protein production rate.

## Results and Discussion

To test our hypothesis, we constructed a translation model consisting of 276 biochemical reactions, including 119 reactions with non-linear kinetics (**Fig. 1**; for details see Methods). This mechanistic model accounts for the concentrations of mRNA, the ribosome, the different charged tRNAs, and the elongation factors Ts (EF-Ts) and Tu (EF-Tu). We fully parameterized the model with molecular masses and kinetic constants measured experimentally ^25–27^; the only exceptions are the initiation parameters, which were previously estimated from gene expression data^25^, and the ribosomal Michaelis constant for the ternary complexes, which was estimated based on the diffusion limit^5^ and hence represents a lower bound. The model is based purely on biochemical and biophysical considerations; it contains no free parameters for fitting, nor does it include any explicit growth-rate dependencies. For *E. coli* growing under different experimental conditions, we used measured growth rates and protein concentrations^28^ to determine the required translation rate and the proportions of the different amino acids incorporated into the elongating proteins. At this required protein production rate, we minimized the combined cost of the translation machinery in our model, treating the concentrations of all components as free variables; the values of individual reaction fluxes result deterministically from these concentrations according to the respective rate laws (**Methods**).

**Figure 1.**
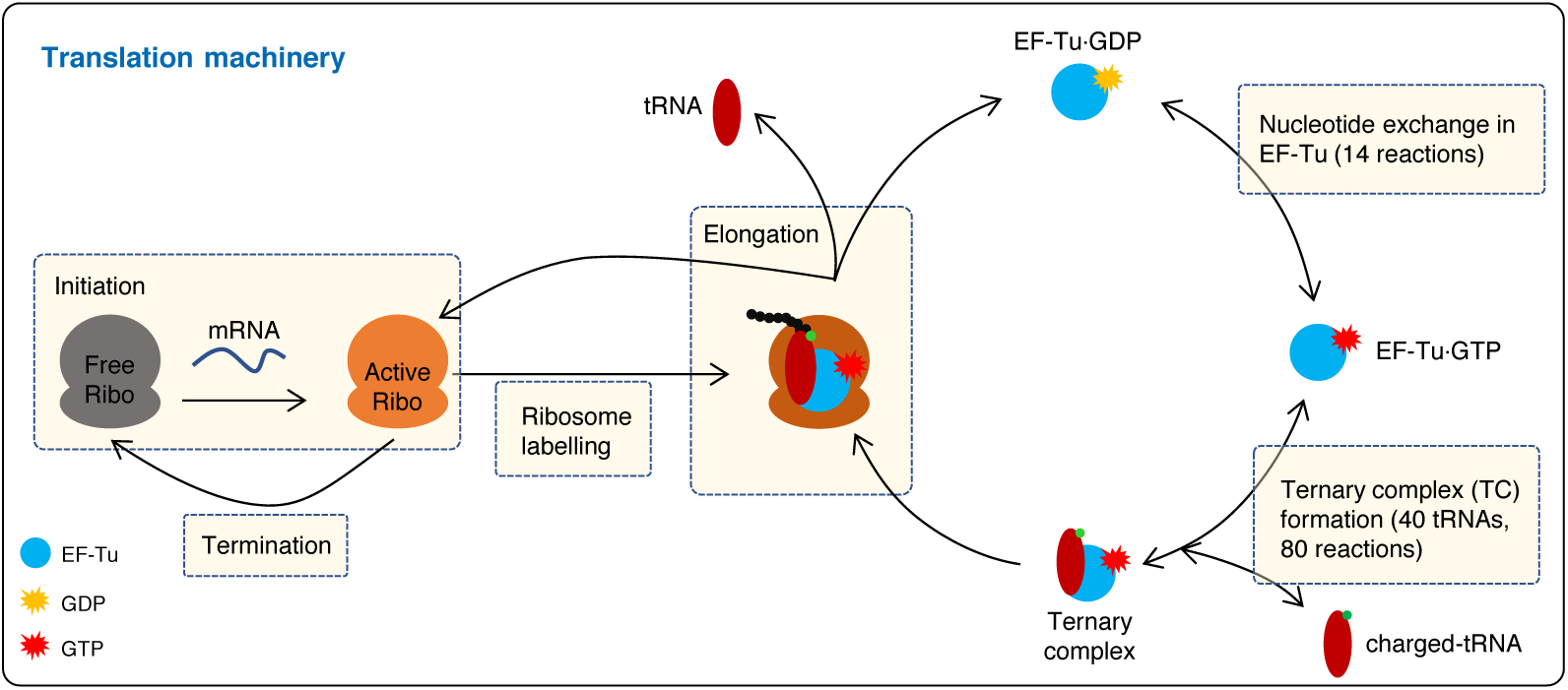
Schematic overview of the translation model. Translation initiation converts the free ribosome to active ribosome by combining it with mRNA. Next, the active ribosome enters elongation, and the codon label is added to limit translation to the cognate ternary complex (TC). The codon-labeled ribosome catalyzes the new peptide bond formation with the TC (EF-Tu·GTP·aa-tRNA) as substrate. EF-Tu·GDP and free tRNA are released after the formation of peptide bond. At the same time with peptide bond formation, the codon labeled ribosome is re-converted to active ribosome, which will be labeled again for the next round of elongation or will go to termination. EF-Tu·GDP is converted to EF-Tu·GTP with the help of EF-Ts. Next, EF-Tu·GTP binds with the charged tRNA (aa-tRNA) to form TC, which is fed into the next round of elongation. Ribosome states are indicated by color: grey=free ribosome; orange= active ribosome; brown=active ribosome with codon label.

We first compared our predictions to experimental data for exponentially growing *E. coli* in different conditions^7–9,28,29^ (see **Fig. 2** for growth in a glucose-limited chemostat at growth rate μ = 0.35 h^-1^; for other conditions, see **Extended Data Fig. 1**). The mechanistic model accurately predicts the absolute concentrations of ribosomes, EF-Tu, EF-Ts, mRNA, and total tRNA in each condition. Predictions for individual tRNA concentrations are less accurate but are still mostly within a 2-fold error (**Fig. 2, Extended Data Fig. 1**); the discrepancies may be due to the simplifying assumption of a single ribosomal Michaelis constant *K*_*m*_ for all tRNA types^5^.

**Figure 2.**
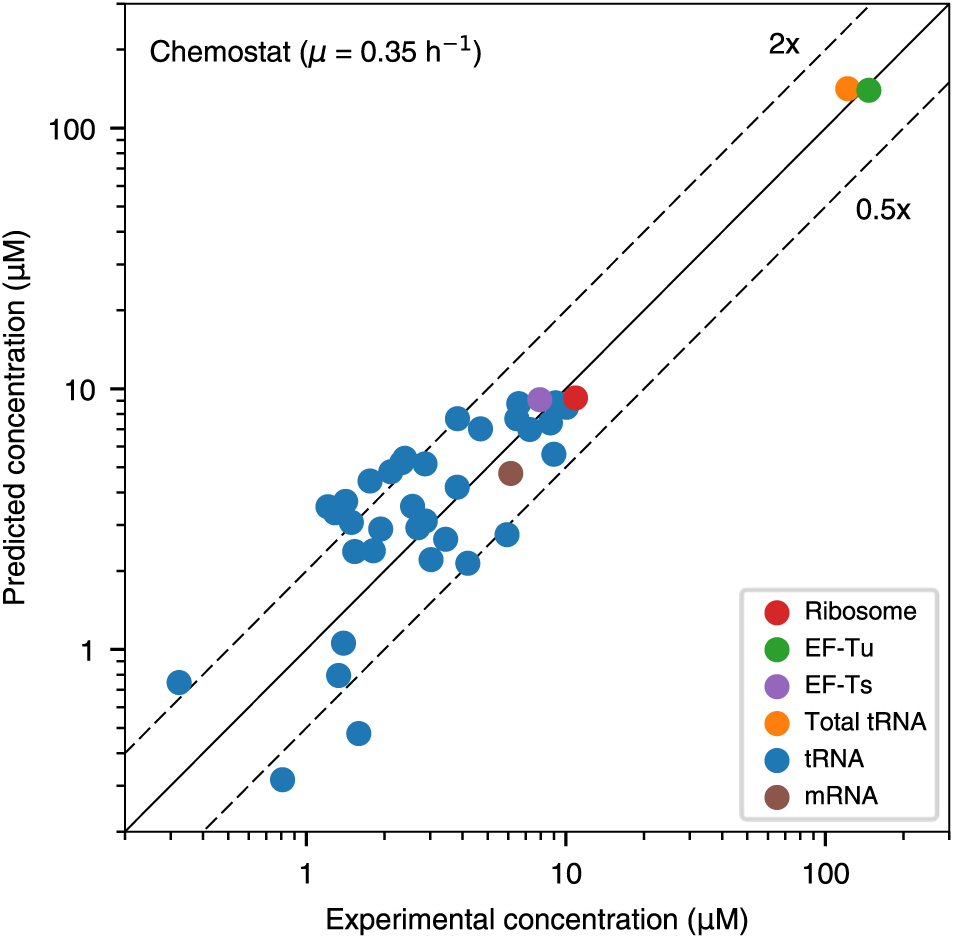
Optimal concentrations of the translation machinery components agree with experimentally measured concentrations in a glucose-limited chemostat (*μ* = 0.35 h^-1^; for other conditions, see **Extended Data Fig. 1**). The solid line shows the expected identity, whereas the upper and lower dashed lines show prediction errors of 2x and 0.5x, respectively. Predictions for ribosome, EF-Tu, EF-Ts, mRNA, and total tRNA are highly accurate, with Pearson’s *R*^2^ = 0.99 and geometric mean fold-error *GMFE* = 1.16, *i*.*e*., predictions based purely on a physico-chemical model and the assumption of cost minimization are on average 16% off. Predictions for individual tRNA species are somewhat less accurate, with *GMFE* = 1.68. Experimentally determined concentrations of the ribosome (averaged over all ribosomal proteins), EF-Tu, and EF-Ts are from Ref. ^28^. mRNA ^29^ and tRNA ^9^ concentrations are interpolated values based on growth rates.

We next tested if this systems-level view on the total cost of translation explains the observed growth rate-dependencies of the expression of translation machinery components^7–9,14,28^, of the elongation rate^14^, and of the RNA/protein ratio^12,14^, considering experimental data across 20 diverse conditions (14 minimal media, including 3 stress conditions; 4 chemostats; and 2 rich media)^28^. The predicted concentrations of ribosomes, EF-Tu, and EF-Ts increase with growth rate in line with experimental observations (**Fig. 3**). At low growth rates (µ<0.3h^-1^; **Fig. 3a**), observed ribosome concentrations exceed those predicted from cost minimization, a deviation consistent with a substantial reserve of deactivated ribosomes at low growth rates^14^. Such deactivated ribosomes may provide fitness benefits in changing environments^18,19^, but cannot be optimally efficient in a constant environment and thus cannot be predicted by our optimization strategy.

**Figure 3.**
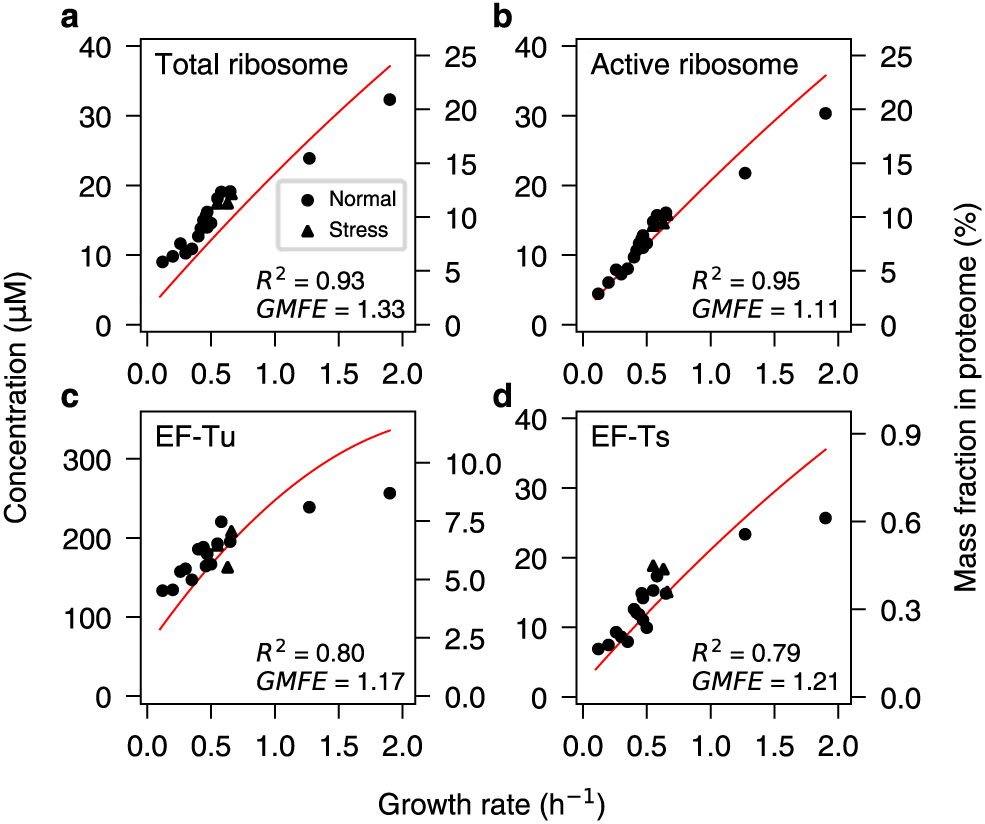
The growth rate dependence of the concentrations of translation machinery components^28^ is consistent between predictions (red lines) and experimental observations. **(a)** Total ribosome concentration (arithmetic means across ribosomal proteins). **(b)** Actively elongating ribosomes, estimated from data in panel (a) according to Ref. ^14^ (see **Methods**), with *R*^2^ = 0.95 and GMFE = 1.11. **(c)** EF-Tu, *R*^2^ = 0.80, GMFE = 1.17. **(d)** EF-Ts, *R*^2^ = 0.79, GMFE = 1.21. Circles indicate normal conditions, triangles indicate stress conditions.

To allow a meaningful comparison between predictions and experiment, we thus estimated the experimental concentration of ribosomes actively involved in elongation (**Methods**). Cost minimization predicts these experimental estimates with high accuracy across the full range of assayed growth rates; observed values deviate from predictions on average by 11% (**Fig. 3b**).

The remaining, non-active ribosome fraction comprises two parts: the deactivated ribosome reserve currently unavailable for translation^14^, and free, potentially active ribosomes not currently bound to mRNA (see **Supplementary Note 2** for the nomenclature on ribosome states). As our model quantifies the abundance of both active and free ribosomes, their subtraction from observed total ribosome concentrations provides an estimate of the deactivated ribosome reserve as a function of growth rate (**Fig. 4**). While this reserve makes up less than 20% of total ribosomes at moderate to fast growth, it reaches almost 50% at the lowest growth rate assayed in Ref.^28^.

**Figure 4.**
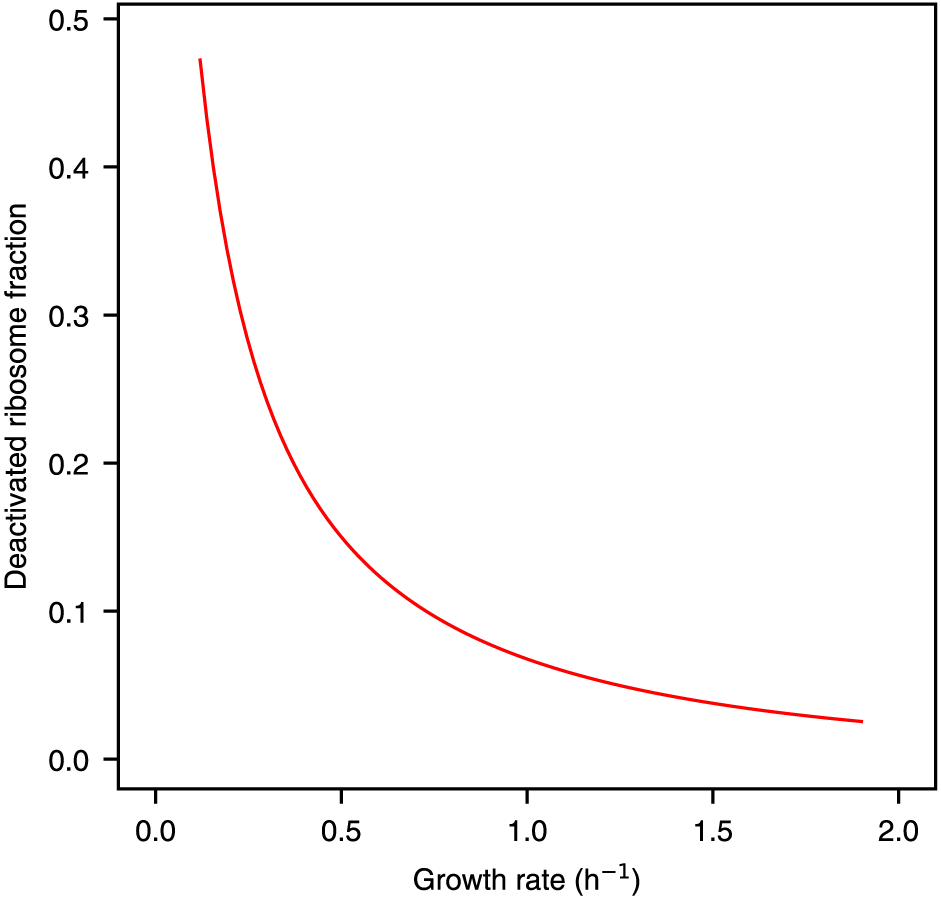
The estimated fraction of deactivated ribosomes increases sharply with decreasing growth rate, reaching almost 50% for the lowest growth rate assayed in Ref. ^28^ and rapidly dropping towards zero at higher growth rates.

The predicted absolute abundances of EF-Tu (**Fig. 3c**), EF-Ts (**Fig. 3d**), and mRNA (**Extended Data Fig. 2a**) also account quantitatively for the experimental data^7–9,28,29^, with average deviations ≤21% in each case. At low growth rates, experimentally observed concentrations of EF-Tu (**Fig. 3c**) and tRNA (**Extended Data Fig. 2b**) are higher than predicted. The model only includes charged (aminoacyl-) tRNA concentrations, and it is likely that the unknown fraction of uncharged tRNA explains at least part of this deviation.

A linear correlation between the RNA/protein ratio and growth rate was discovered in the 1950s^3,20,21,30^ and forms the basis of phenomenological bacterial growth laws^5,12,14^. Relating the predicted total RNA (ribosomal RNA + tRNA + mRNA) with measured protein concentrations^28^ indeed results in a near-linear relationship, accurately matching observed values at high to intermediate growth rates (µ > 0.3 h^-1^; **Fig. 5a**). At lower growth rates, model predictions are slightly too low, likely because of the deactivated ribosome reserve^14^ (**Fig. 4**). At low growth rates (µ = 0.12 h^-1^), RNA and proteins allocated to an optimally efficient translation machinery (including deactivated ribosomes) account for 12% of total dry mass, rising almost linearly to ∼45% at high growth rates (µ = 1.9 h^-1^; **Extended Data Fig. 3**).

**Figure 5.**
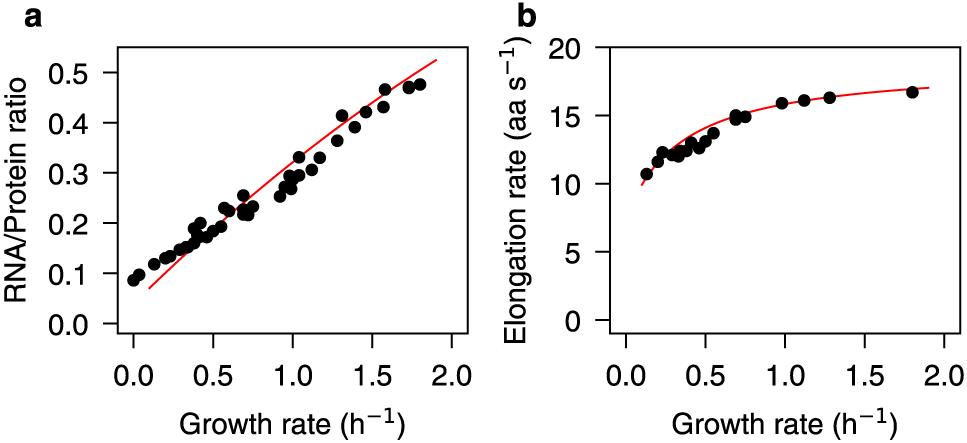
The growth rate dependences of the total RNA/protein ratio and ribosome activity are consequences of translation machinery cost minimization. **(a)** Predicted total RNA concentration (mRNA + tRNA + rRNA) relative to observed total protein concentration at different cellular growth rates (red line) compared to experimental observations^12,14^; *R*^2^ = 0.97, *GMFE* = 1.10. **(b)** Predicted (red line) and experimentally determined^14^ elongation rates of actively translating ribosomes (ribosome activities); *R*^2^ = 0.93, *GMFE* = 1.03. At the lowest assayed growth rates, non-growth related translation – which is not included in the model – may become comparable to growth-related translation; at these growth rates, the numerical optimization of our model did not converge, and thus the red lines are not extended into this region.

The concentrations of the individual components of the translation machinery determine the average translation elongation rate (ribosomal activity), defined as the total cellular translation rate divided by the total active ribosome content^19^. The predicted elongation rates closely match the experimental data^14^ over a broad range of growth rates (**Fig. 5b**).

The expression of *E. coli*’s translation machinery reacts strongly to the exposure to antibiotics that inhibit the ribosome, such as chloramphenicol^12,14,15^. The details of these changes can also be understood from our hypothesis of cost minimization. The concentrations of ribosomes and EF-Tu, the RNA/protein ratio, and the elongation rate of active ribosomes increase under chloramphenicol stress (**Extended Data Fig. 4**); these changes partially compensate for the reduced fraction of active ribosomes. The concentration of EF-Ts instead decreases with increasing chloramphenicol concentration (**Extended Data Fig. 4c**). EF-Ts contributes to translation by converting EF-Tu·GDP to EF-Tu·GTP, which then forms a ternary complex with charged tRNA. Under chloramphenicol stress, fewer ternary complexes are turned over, and hence less EF-Ts is needed.

In sum, cost minimization in a mechanistic bottom-up model of optimal translation efficiency, fully parameterized with known kinetic constants and molecular masses, accounts quantitatively for the concentrations of all molecule species involved. The optimal concentrations of different components change differentially with growth rate, explaining the observed scaling of *E. coli*’s translation machinery composition, RNA composition, and elongation rate. We conclude that *E. coli*’s translation machinery works close to optimal efficiency in terms of the fraction of total dry mass it occupies. This fraction comprises all molecule species involved in translation, not only the protein part as suggested earlier^5,31^. Our results further support the idea that phenomenological growth laws of proteome composition^5,12,14,15^ may have their root in the costs associated with the non-protein molecules involved in particular processes, and that their explicit inclusion in systems biology models of cellular growth^5,25,32,33^ may eventually allow these models to abandon any reliance on phenomenological parameters.

## Acknowledgments

This work was supported by the German Research Foundation (DFG grants IRTG 1515 supporting HD; EXC 1028, CRC 680, and CRC 1310 to MJL). We thank Matteo Mori and Terrence Hwa for helpful discussions.

## Endnotes

### Author Contributions

HD conceived of the study. XPH developed and implemented the model and performed the analyses. MJL supervised the study. XPH and MJL interpreted the results and wrote the manuscript.

The authors declare that they have no competing interests.

Correspondence and requests for materials should be addressed to.

## Extended Data Figures

**Extended Data Figure 1.**
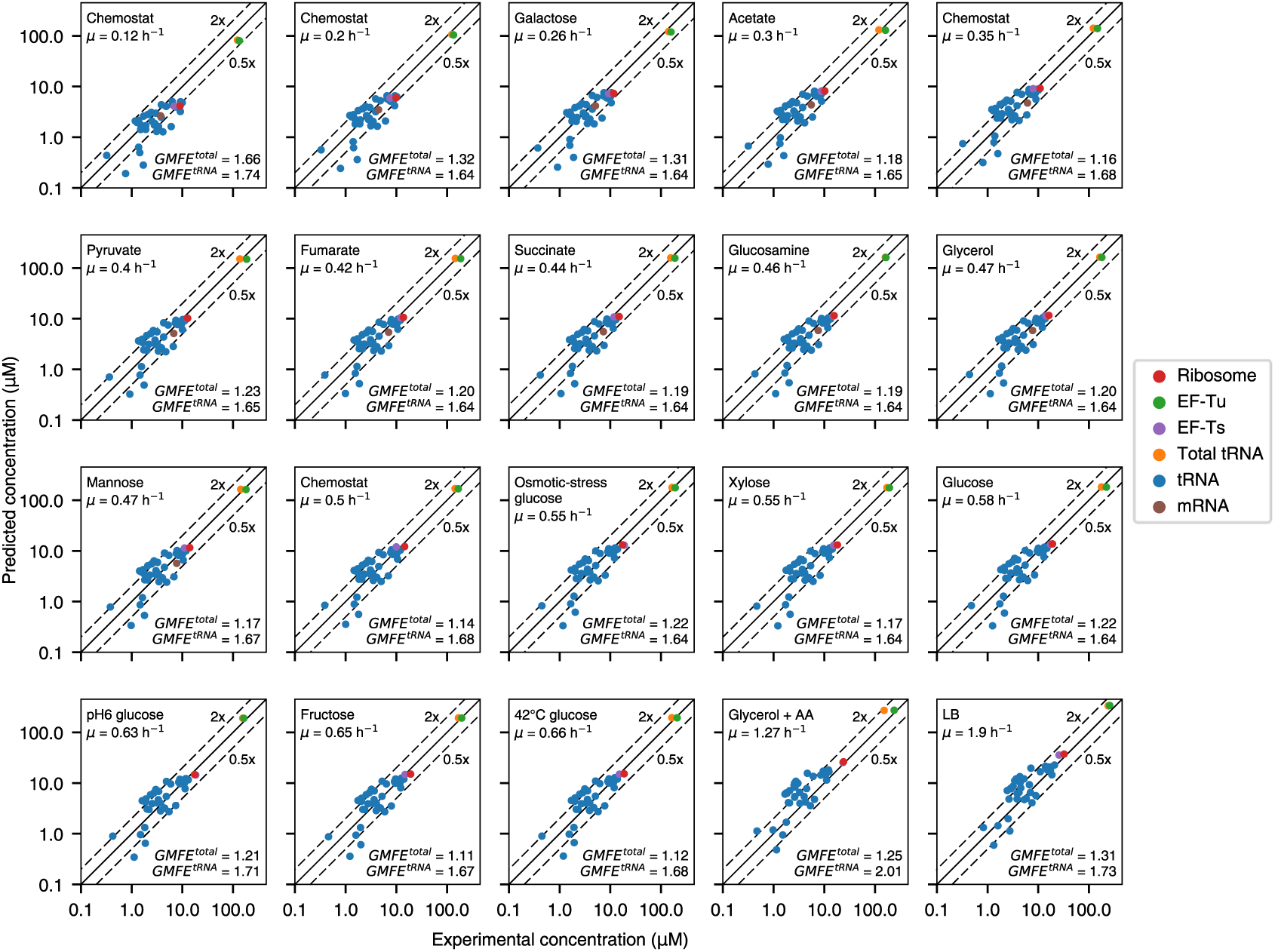
Translation machinery at optimal state at 20 growth conditions on different media and in chemostats with a minimal glucose medium, sorted by ascending growth rate. The conditions are those under which protein concentrations were measured in Ref. ^28^. mRNA ^29^ and tRNA ^9^ were assayed at different growth rates; in order to compare all data at the same growth rates, we chose the growth rate at which protein concentrations (ribosome, EF-Tu, EF-Ts) were measured as the reference and used quadratic regression models across the available data to estimate corresponding mRNA and tRNA concentrations. As absolute mRNA concentration is only available from low to intermediate growth rates (0.11 to 0.49)^29^, we did not attempt to infer mRNA concentration outside of this range.

**Extended Data Figure 2.**
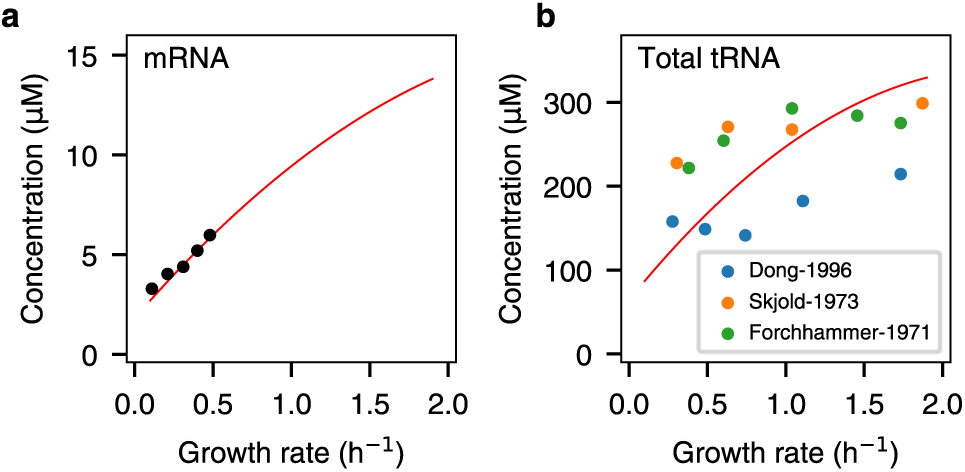
The concentrations of the major non-ribosomal RNA pools predicted from cost minimization are consistent with experimental observations. **(a)** mRNA ^29^, *R*^2^ = 0.97, GMFE = 1.06. **(b)** Total tRNA data from Dong *et al*. ^9^ (summed over individual tRNAs), Forchhammer *et al*. ^8^, and Skjold *et al*. ^7^; combined *R*^2^ = 0.27, *GMFE* = 1.30.

**Extended Data Figure 3.**
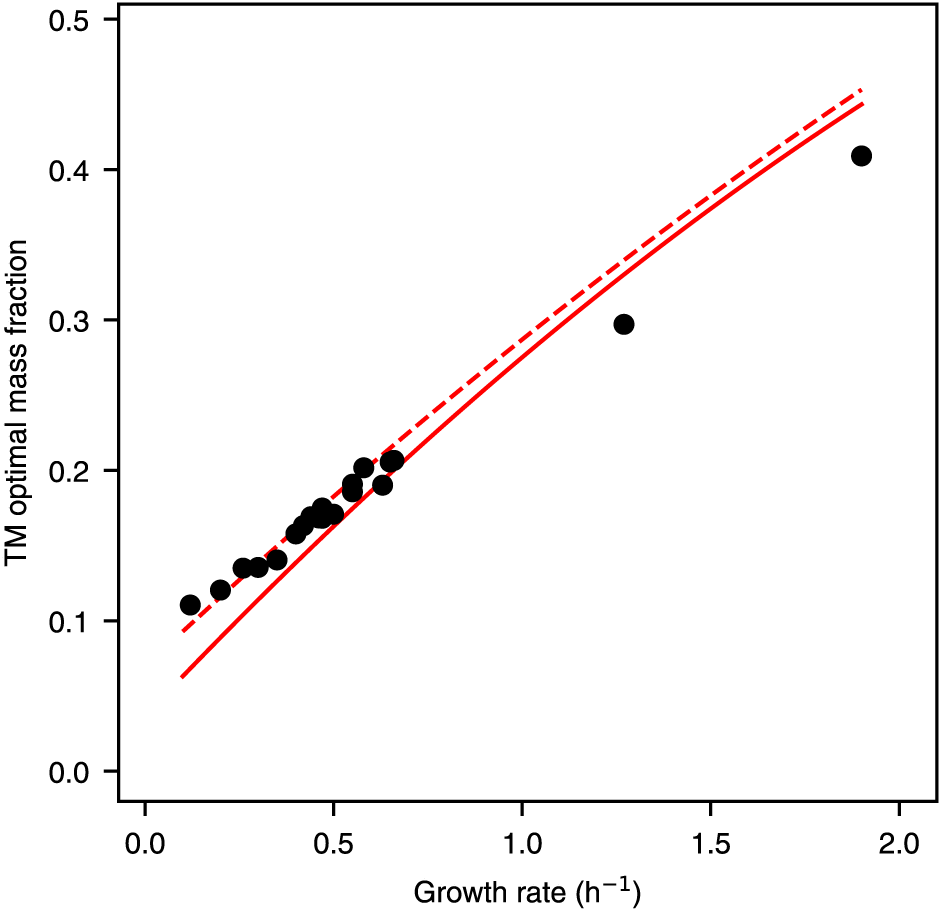
Theoretically optimal resource allocation to the translation machinery as a fraction of total dry mass increases almost linearly with growth rate. The solid red line indicates the model predictions, without accounting for deactivated ribosomes. The dashed line indicates the predicted optimal mass fraction when we additionally include the fraction of deactivated ribosomes, which cannot be predicted by a steady-state model but which we estimated from experimental observations (see Methods for details). Experimental data (points) sums the observed concentrations of translation associated proteins^28^ (ribosomal proteins, EF-Tu, EF-Ts) and RNA^12,14^ (ribosomal RNA, tRAN, mRNA; interpolated to the same growth rates as in the protein measurements, see Methods). Note that the mass fraction of the translation machinery does not include GDP, GTP, free tRNA, tRNA-synthetases, and elongation factor G (*fusA*). Mass fractions are calculated based on the assumption of a constant proteome mass fraction of 50% of the total dry mass. Some experimental data shows that the mass fraction of protein in total dry weight decreases slightly with growth rate^19,29^, and thus at high growth rates the translation machinery mass fraction may be slightly lower than shown.

**Extended Data Figure 4.**
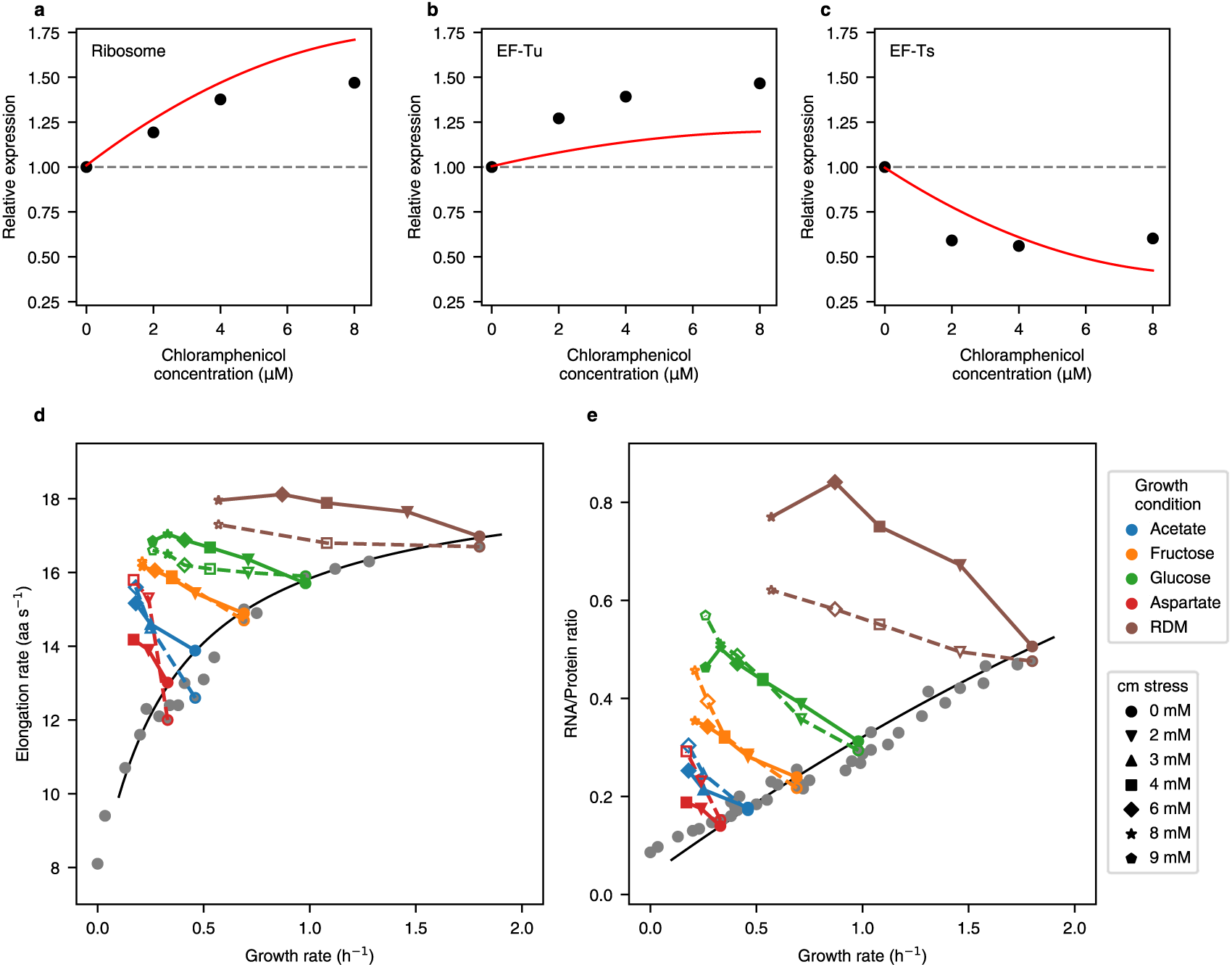
Optimality of the translation machinery under chloramphenicol stress. Model predictions (red lines) of relative changes in the concentrations of **(a)** ribosome, **(b)** EF-Tu, and **(c)** EF-Ts under increasing chloramphenicol stress are qualitatively consistent with experimental data^15^. Predicted **(d)** elongation rates and **(e)** RNA/protein ratios under chloramphenicol stress are also qualitatively consistent with experimental data^14^. Grey dots indicate experimental elongation rates without chloramphenicol stress; the black line marks the corresponding (non-stressed) predictions. Different symbols indicate varying chloramphenicol concentrations, while colours indicate growth conditions (different nutrients). Dashed lines connect experimental elongation rates (open symbols) under chloramphenicol stress on the same nutrient; solid lines connect the corresponding elongation rate predictions (filled symbols). Chloramphenicol concentrations were varied from 0mM to 9mM. In both predictions and experiment, elongation rates increase with growing chloramphenicol stress, with faster increases under progressively poorer nutrient conditions. The overestimated RNA/protein ratio on rich defined medium (RDM) likely reflects the fact that ribosome is inhibited less by chloramphenicol *in vivo* than theoretical calculations predict (see Fig. N1 in Ref.^14^). The predictions are functions of the growth rate and of chloramphenicol concentration; the non-smoothness of the prediction lines likely arise from experimental uncertainties in the corresponding values.

