## Supplementary Information for "The protein translation machinery is expressed for maximal efficiency in *Escherichia coli*"

### Methods

#### Experimental concentrations of all components

We used molar concentrations (µM) in the model; thus, all experimental data were converted to molar concentrations.

***Ribosomes, EF-Tu, and EF-Ts.*** We used (1) protein copy number per cell[1] and (2) cell volume[2] to calculate the molar concentrations of ribosomes, EF-Tu, and EF-Ts as

$$c_i = \frac{N_i}{V_{\text{cell}} N_{\text{A}}}, \qquad (1)$$

where $N_i$ is the protein copy number per cell of the considered molecule species, $V_{\text{cell}}$ is the cell volume at the given condition, and $N_{\text{A}}$ is the Avogadro constant. We estimated the ribosome concentration as the arithmetic mean concentration of all ribosomal proteins.

In a more recent publication[3], the authors of Ref. [2] re-measured the volume of cells by super-resolution microscopy and found that cell volume was overestimated in Ref. [2] (by a factor of 1/0.67 for growth on glucose). Given that the absolute protein concentration was assumed to be constant across all conditions and glucose was chosen as the reference to calculate absolute protein concentrations (copy numbers per cell) [1], we thus modified cell volume by a factor of 0.67 relative to the values in Ref. [2] for all conditions.

***Active ribosome.*** Active ribosomes are defined here as ribosomes engaged in peptide elongation. Dai et al. estimated the fraction of active ribosomes, $f_{\text{active}}$, in *E. coli* at different growth rates[4]. We fitted a Michaelis-Menten type equation to their data, resulting in $f_{\text{active}} = \mu / (0.124 + \mu)$. For each total ribosome concentration $c_{\text{ribosome}}$ in Ref. [1], we then estimated the corresponding active ribosome concentration as $c_{\text{active-ribosome}} = f_{\text{active}} * c_{\text{ribosome}}$.

***GTP and GDP.*** GTP and GDP concentrations are from Ref. [5]. We chose the data for growth on glucose for all simulations (See **Supplementary Note 3** for details).

***mRNA.*** mRNA concentration was calculated from the data given in from Ref. [6] as the ratio of mRNA copy number per cell and cell volume. To estimate the mRNA concentrations at the growth rates shown in Fig. 1 and Extended Data Fig. 1, we fitted the mRNA concentration as a quadratic function of growth rate and read off the values at the required growth rates. mRNA concentrations were assayed only at growth rates between 0.11 $h^{-1}$ and 0.49 $h^{-1}$, and we did not attempt to extrapolate values beyond this range.

***tRNA.*** We collected three independent datasets of tRNA concentrations. Dataset1[7] contains tRNA concentration for each individual tRNA, whereas both dataset2[8] and dataset3[9] contain only total tRNA concentrations. In all datasets, tRNA abundance was measured as the ratio of tRNA to ribosomal RNA (rRNA). We scaled these values to absolute tRNA concentrations assuming that the rRNA concentration corresponds to the ribosome concentration estimated from the proteomics data (see the subsection "Ribosome, EF-Tu, and EF-Ts").

***Concentrations of individual tRNAs and relationship to our model.*** Our model differentiates tRNAs by their anticodons (see Model for details). Thus, 40 tRNAs were used to represent all elongator tRNAs. The tRNAs modeled in this work are listed in Table S2 together with their common names used in dataset1[7] and their gene IDs.

In the experiments by Dong et al. (dataset1)[7], tRNAs were classified into 41 distinct sets based on two-dimensional polyacrylamide gel electrophoresis. We combined two tRNA sets corresponding to different tRNA weights if they have the same anticodon (*i.e.*, the pairs of Val2A + Val2B, Thr1 + Thr3, and Tyr1 + Tyr2). The experimenters could not distinguish between the tRNAs Gly1 and Gly2, as these have very similar molecular weights and isoelectric point; the same was true for Ile1 and Ile2. We estimated the individual concentrations of these four tRNAs based on the ratios 3:2 between Gly1 and Gly2 and 20:1 between Ile1 and Ile2 observed by Ikemura *et al.* [10].

To estimate the tRNA concentrations at the growth rates shown in Fig. 2 and Extended Data Fig. 1, we fitted the concentration of each tRNA in dataset1 (measured for growth rates ranging from 0.28 $h^{-1}$ to 1.73 $h^{-1}$) to a quadratic function of growth rate and extended this function to the required range, [0.12 to 1.9].

***Combination of tRNAs that are predicted to be non-expressed with their co-functional tRNAs.*** The predicted concentrations of 6 tRNAs are 0 μM; these are highlighted in red in Table S4. This result is a straightforward consequence of the model structure. The relationship between codons and tRNAs is not one-to-one in *E. coli*. Consider a given codon (codon1) that has more than one cognate tRNA, labeled tRNA1 and tRNA2. If tRNA2 is also the cognate tRNA of another codon (codon2), the predicted concentration of tRNA1 will be zero: for the same "price" (the same contribution to the limited total mass concentration), tRNA2 can service two codons, while tRNA1 can service only one. For example, codon GGG has two cognate tRNAs, gly1 and gly2; gly2 is also the cognate tRNA of codon GGA. Thus, both gly1 and gly2 can translate GGG, but gly2 can translate GGA, too, and is thus more valuable to the cell if we assume that both tRNAs are processed equally efficiently, as done in the model. Therefore, the predicted optimal concentration of gly1 will be zero (note that this might not occur in models that consider different ribosomal $k_{cat}$ or $K_m$ values for the two tRNAs). To compare our predictions to the experimental data[7], we combined tRNAs with predicted zero concentration with their co-functioning tRNAs in both the predictions and the experimental data. The resulting six combined tRNA pairs are: GLy1 + Gly2; Leu1 + Leu3; Leu4 + Leu5; Pro1 + Pro3; Ser2 + Ser1; Thr2 + Thr4. In all reported figures, the total number of tRNAs shown is thus 34.

***Concentration of total protein ($c_{protein}$) and amino acids encoded by each codon ($c_{codon\text{-}i}$).*** Let $P$ be the set of all proteins quantified experimentally under a specific growth condition. Then

$$c_{\text{protein}} = \frac{\sum_{p \in P} N_p}{V_{\text{cell}} N_{\text{A}}} \tag{17}$$

is the total molar protein concentration, where $N_p$ is the abundance (copy number) of protein $p$ in one cell, $V_{\text{cell}}$ is the cell volume, and $N_{\text{A}}$ is the Avogadro constant.

The concentration of amino acids encoded by *codon-i* is calculated from genome-scale protein expression data, cell volume, and the mRNA sequences:

$$c_{codon-i} = \frac{\sum_{p \in P} N_p N_{ip}}{V_{\text{cell}} N_{\text{A}}}, \tag{18}$$

with $N_{ip}$ the number of *codon-i* occurrences in the mRNA encoding protein $p$.

For simulations under chloramphenicol stress, $c_{\text{protein}}$ and $c_{codon\text{-}i}$ are not available. We approximated their values by the corresponding concentrations for growth on glucose in the absence of the antibiotic.

#### Mass fraction of the translation machinery in total dry weight (Extended Data Fig. 3)

The mass fraction of the translation machinery estimated in Extended Data Fig. 3 includes ribosome, mRNA, charged-tRNA, EF-Tu, and EF-Ts and does not include GDP, GTP, free tRNA, tRNA-synthetases, and elongation factor G (*fusA*). We converted from protein fractions to mass fractions of total dry weight assuming that the mass fraction of protein in total dry weight was 50% for all growth rates[1].

***Experimental estimate.*** The mass fraction of the translation machinery in total dry weight is the sum of two parts: (1) protein and (2) RNA.

(1) We calculated the mass fraction of translational proteins (including ribosomal protein, EF-Tu, and EF-Ts) in dry weight from the proteomics data in Ref. [1].

(2) We fitted the reported total RNA/protein ratio in Refs.[4,11] to a quadratic function of growth rate. We then used this fitted function to calculate the RNA/protein ratio at the growth rates assayed by Schmidt *et al.* [1].

***Theoretical prediction.*** The predicted mass fraction of the translation machinery was the ratio of the total mass concentration of the translation machinery (including free ribosome, active ribosome, EF-Tu, EF-Ts, aa-tRNA, and mRNA) to the total dry mass density (assumed to be twice the synthesized protein density). For the prediction including de-activated ribosome concentrations (dashed line in Extended Data Fig. 3), we added estimates of de-activated ribosome concentrations according to Fig. 4 of the main text.

#### Molecular weights

Molecular weights of ribosome, charged tRNAs (aa-tRNAs), EF-Tu, EF-Ts, and the ternary complexes (TC, EF-Tu·GTP·aa-tRNA) were calculated from their sequences. The stoichiometry of ribosomal proteins and RNAs in the ribosome was obtained from the EcoCyc database[12]; the stoichiometry of all components is 1 except for RplL, for which it is 4.

We used an average mRNA to represent the total mRNA. The molecular weight of an average mRNA ($MW_{\text{mRNA}}$) is the sum of two parts: (1) the molecular weight of the coding sequence (CDS) of mRNA ($MW_{\text{mRNA-CDS}}$), which was calculated from protein-expression-weighted mRNA length and nucleotide composition of *E. coli* protein-coding sequences in each growth condition[1]; (2) the weight of the untranslated region (UTR) of mRNA ($MW_{\text{mRNA-UTR}}$), which was calculated from the average nucleotide composition of the genome and the typical length of UTR. The length of UTR was assumed to be 85 nt, a typical length of the untranslated region in *E. coli*[13]. Thus,

$$MW_{\text{mRNA}} = MW_{\text{mRNA}-\text{CDS}} + MW_{\text{mRNA}-\text{UTR}}. \quad (2)$$

In the simulations of translation under antibiotic stress, the molecular weight of chloramphenicol was set to 0.

#### Data availability

The optimization problem, including the model and its parameterization, is provided as a GAMS file.

### Model

The mechanistic translation model encompasses the processes of translation initiation, elongation, termination, nucleotide exchange in EF-Tu, and ternary complex (TC) formation, illustrated in Fig. 1 of the main text. In total, the model includes 276 reactions.

#### Initiation

During initiation, mRNA converts free ribosomes to active ribosomes:

$$\text{Ribosome}_{\text{free}} \rightarrow \text{Ribosome}_{\text{active}}. \tag{r1}$$

The initiation rate is modelled with Michaelis–Menten kinetics, where mRNA acts as the “enzyme” (with concentration $c_{\mathrm{mRNA}}$) and free (unbound) ribosomes act as the substrate[14] (with concentration $c_{\mathrm{ribo-free}}$),

$$v_{\mathrm{tl-init}} = k_{\mathrm{cat-mRNA}} \cdot c_{\mathrm{mRNA}} \cdot \frac{c_{\mathrm{ribo-free}}}{K_{\mathrm{M-ribo}} + c_{\mathrm{ribo-free}}}, \tag{3}$$

with $k_{\text{cat–mRNA}}$ = 1.33 $s^{-1}$ and $K_{\text{M-ribo}}$ = 8.5 µM from Ref. [14]. We assume that total mRNA concentration is much greater than free ribosome concentration, and we hence do not distinguish free and total mRNA.

We assume that the turnover number of mRNA for ribosome binding ($k_{\mathrm{cat-mRNA}}$) is growth rate-independent, and hence that the observed growth rate-dependent activity is due to changes in the concentration of free ribosomes available for initiation[15]. Thus, we used the maximal reported mRNA activity as an estimate of $k_{\mathrm{cat-mRNA}}$.

#### Ternary complex formation and nucleotide exchange in EF-Tu

Ternary complex formation and nucleotide exchange in EF-Tu are the processes by which the translation machinery recycles its substrates, the ternary complexes, for elongation.

***TC formation.*** The binding of EF-Tu·GTP to charged-tRNA (aa-tRNA) forms the ternary complex (reaction 8 in Figure S1), which is the substrate of translation elongation.

***Nucleotide exchange in EF-Tu.*** EF-Tu·GDP is released after the formation of a new peptide bond. Elongation factor Ts (EF-Ts) binds to EF-Tu·GDP and induces the exchange of GDP for GTP (reactions 1 to 7 in Figure S1).

The individual steps of these two processes are modelled with mass action kinetics[16,17] (Figure S1).

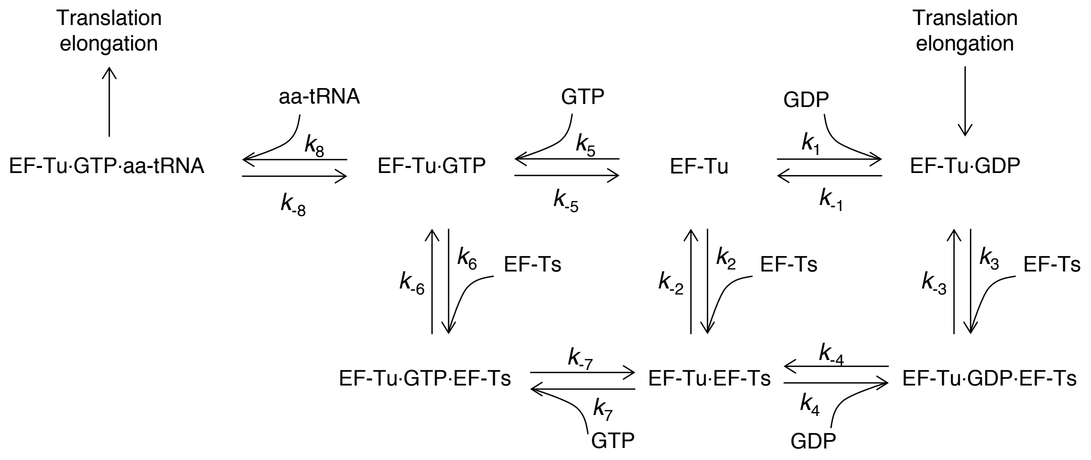


**Figure S1.** The process of nucleotide exchange in EF-Tu (reactions 1 to 7) and ternary complex (TC) formation (reaction 8). The $k_i$ symbols represent the mass action kinetic rate constants, with reaction rates $v_i = k_i \prod[c]$ [16,17].

In the implementation of the model, we divide each reversible reaction into two irreversible reactions. The TC formation reaction (reaction 8) is a set of reactions that include the binding of 40 aa-tRNAs to Tu·GTP, and its rate constants $k_8$ and $k_{-8}$ depend on which amino acid is involved[17] (for the parameter values see Table S1.)

#### Elongation

Elongation is a very complex process[18,19]. For simplicity, we model elongation as a single reaction with an active ribosome as the enzyme and TC as the substrate, which was proposed by *Klumpp et al.*[20]. In this single reaction model, TC is discriminated by the anticodon and all TCs are treated with the same activity. Michaelis–Menten kinetics are used to describe the reaction rate[20]. There are 40 anticodons in total for all elongator tRNAs in *E. coli*; accordingly, our model uses 40 tRNAs to represent all tRNAs.

At steady state, the total translation rate (per cytosolic volume) of each codon remains constant. We do not model the translation of a whole protein. Instead, we decompose protein synthesis into the translation of 61 codons (see also "Modeling" below).

***Active ribosome state***

We distinguish active ribosomes according to their binding codons (codon in ribosome A site). Thus, there are 61 types of active ribosome in the model, distinguished by codon labels:

$$\text{Ribosome}_{\text{active}} \rightarrow \text{Ribosome}_{\text{codon-i}}, \qquad \text{(r2)}$$

where *codon-i* is one of the 61 codons and $Ribosome_{codon\text{-}i}$ is the active ribosome that binds *codon-i*. This reaction is constrained by mass balance, but is considered to be instantaneous. We assume that when an active ribosome binds with a specific codon, it only translates the codon's cognate tRNA.

In the model, there are 61 codons and 40 tRNAs, and the relation between tRNA and codon is not one-to-one. Based on the number of cognate tRNAs, we partition the 61 codons into 2 classes: class 1 codons have one cognate tRNA, whereas class 2 codons have two cognate tRNAs.

For class 1 codons ($n$ = 51), the elongation reaction is:

$$\mathrm{TC_{codon\text{-}i}} \xrightarrow{\mathrm{Ribosome_{codon\text{-}i}}} \mathrm{EF\text{-}Tu{\cdot}GDP+tRNA_{codon\text{-}i}+aa_{codon\text{-}i}}, \tag{r3}$$

where $TC_{codon\text{-}i}$ is the cognate TC of *codon-i*, $tRNA_{codon\text{-}i}$ is the released free tRNA, and $aa_{codon\text{-}i}$ symbolizes the amino acid that was just appended to the growing peptide. Simultaneously, the $Ribosome_{codon\text{-}i}$ is translocated to the next round of elongation,

$$\mathrm{Ribosome_{codon\text{-}i}} \rightarrow \mathrm{Ribosome_{active}}. \tag{r4}$$

For simplicity, we combine r3 and r4 as the new reaction r5, with $Ribosome_{codon\text{-}i}$ as a substrate and $Ribosome_{\mathrm{active}}$ as a product (the same for r6 and r7),

$$\mathrm{TC_{codon\text{-}i}} + \mathrm{Ribosome_{codon\text{-}i}} \xrightarrow{\mathrm{Ribosome_{codon\text{-}i}}} \mathrm{Ribosome_{active}} + \mathrm{EF\text{-}Tu{\cdot}GDP+tRNA_{codon\text{-}i}+aa_{codon\text{-}i}}. \tag{r5}$$

The translation rate of *codon-i* is described by Michaelis-Menten kinetics,

$$v_{tl-codon-i} = c_{ribo-codon-i} \cdot k_{\mathrm{cat-ribo}} \cdot \frac{c_{TC-codon-i}}{c_{TC-codon-i} + K_{\mathrm{M-TC}}}, \tag{4}$$

where $c_{ribo\text{-}codon\text{-}i}$ is the concentration of ribosome that binding with *codon-i* ($Ribosome_{codon\text{-}i}$), $c_{TC\text{-}codon\text{-}i}$ is the concentration of cognate TC of *codon-i* ($TC_{codon\text{-}i}$), $k_{\mathrm{cat\text{-}ribo}} = 22\ \mathrm{s}^{-1}$ , and $K_{\mathrm{M\text{-}TC}} = 3\ \mu\mathrm{M}$ (parameters from Ref. [20]).

For class 2 codons ($n$ = 10), the active ribosome can translate two TCs and thus there are two reactions:

$$\mathrm{TC_{codon\text{-}i\text{-}1}} + \mathrm{Ribosome_{codon\text{-}i}} \xrightarrow{\mathrm{Ribosome_{codon\text{-}i}}} \mathrm{Ribosome_{active}} + \mathrm{EF\text{-}Tu{\cdot}GDP+tRNA_{codon\text{-}i\text{-}1}+aa_{codon\text{-}i}} \tag{r6}$$

and

$$\mathrm{TC_{codon\text{-}i\text{-}2}} + \mathrm{Ribosome_{codon\text{-}i}} \xrightarrow{\mathrm{Ribosome_{codon\text{-}i}}} \mathrm{Ribosome_{active}} + \mathrm{EF\text{-}Tu{\cdot}GDP+tRNA_{codon\text{-}i\text{-}2}+aa_{codon\text{-}i}}. \tag{r7}$$

The translation rate of *codon-i* is the sum of these two reactions:

$$v_{tl-codon-i} = v_1 + v_2, \tag{5}$$

with

$$v_1 = c_{ribo-codon-i} \cdot k_{\mathrm{cat-ribo}} \cdot \frac{c_{TC-codon-i-1}}{(c_{TC-codon-i-1} + c_{TC-codon-i-2}) + K_{\mathrm{M-TC}}} \tag{6}$$

and

$$v_2 = c_{ribo-codon-i} \cdot k_{\mathrm{cat-ribo}} \cdot \frac{c_{TC-codon-i-2}}{(c_{TC-codon-i-1} + c_{TC-codon-i-2}) + K_{\mathrm{M-TC}}}. \tag{7}$$

#### Termination

In termination, an active ribosome is converted to a free ribosome:

$$\text{Ribosome}_{\text{active}} \rightarrow \text{Ribosome}_{\text{free}}. \tag{r8}$$

The termination rate is equal to the protein synthesis rate at steady state:

$$v_{\text{term}} = v_{\text{protein-syn}} = \mu \cdot c_{\text{protein}}, \tag{8}$$

where $c_{\text{protein}}$ is the absolute concentration of protein measured experimentally.

#### Exchange reactions

Besides the reactions mentioned above, we also add exchange reactions that allow the influx of free ribosome, charged tRNA, mRNA, EF-Tu, EF-Ts, GTP, and GDP into the system. We also add exchange reactions that allow efflux of free tRNA and GDP out of the system.

#### Antibiotic stress

To model chloramphenicol (cm) stress, we add the exchange reaction for chloramphenicol. All forms of ribosome can be inhibited by chloramphenicol:

$$\text{Ribosome+cm} \leftrightarrow \text{Ribosome} \cdot \text{cm}, \tag{r9}$$

where Ribosome includes both free and active ribosomes. The reaction rate is described by mass action kinetics with $k_{\text{on}}$ = 0.00057 μM·$s^{-1}$ (0.034 μM·$\text{min}^{-1}$) and $k_{\text{off}}$ = 0.0014 $s^{-1}$ (0.084 $\text{min}^{-1}$)[21].

For simplicity, we make the following assumptions:

(1) chloramphenicol diffuses freely across cell membranes, such that the intracellular concentration of free chloramphenicol is the same as that in the medium;
(2) chloramphenicol bound to an active ribosome (ribosome·cm) causes the ribosome·cm complex to dissociate quickly from the mRNA, thus not affecting further translation of the mRNA.

We assume that the reaction of ribosome and chloramphenicol binding is at steady state (dynamic equilibrium), and thus the active ribosome concentration will be constant. However, not all active ribosomes will be able to successfully finish translation, as the active ribosome can be inhibited by chloramphenicol during translation. Thus, to estimate the production rate of functional proteins, we need to estimate the probability that the active ribosome can finish translation without chloramphenicol inhibition.

To calculate this probability, we use the method proposed in Ref. [4]. The probability of chloramphenicol binding to a ribosome in a given time unit is:

$$k_{\text{hit}} = k_{\text{on}} c_{\text{cm}}, \tag{9}$$

where $k_{\text{on}}$ is the binding constant. The probability that the ribosome is bound $n$ times in the time interval $t$ follows the Poisson distribution

$$P(n) = e^{-k_{\text{hit}} t_{\text{tl}}} \frac{k_{\text{hit}} t_{\text{tl}}}{n!}, \tag{10}$$

where $t_{\text{tl}}$ is the experimental measured translation time of the translated gene. The probability that the ribosome can finish translation without inhibition by chloramphenicol is $P(0)$:

$$P_{\text{tl}} = e^{-k_{\text{hit}} t_{\text{tl}}}. \tag{11}$$

Here, $t_{\text{tl}}$ is the time for LacZ and $t_{\text{tl}}$ = 72 s (from Ref. [4]). For simplicity, we assume that all codon-labeled active ribosomes (61 forms of active ribosome) have the same $P_{\text{tl}}$, and thus the effective concentration of *codon-i* labeled ribosome is

$$c_{ribo-eff-i} = P_{\text{tl}} c_{ribo-i}. \tag{12}$$

Under inhibition by chloramphenicol, this effective ribosome concentration of *codon-i* ($c_{ribo\text{-}eff\text{-}i}$) replaces the ribosome concentration of ($c_{ribo\text{-}i}$) in the model.

#### Modeling

We assume translation at steady state and use a constraint-based optimization model. The constraints are given by the above equations and by the requirement of a given total cellular rate of protein synthesis (estimated as the product of growth rate and experimental proteome composition at this growth rate). For simplicity, we decomposed protein synthesis into the translation of 61 codons, and so the constraint on protein synthesis rate is implemented as 61 individual equations, each representing the translation rate of one codon. At steady state, the codon translation rate equals the dilution rate of synthesized amino acids encoded by *codon-i*, i.e., the growth rate $\mu$ multiplied by the concentration of amino acids coded by *codon-i* in proteome data:

$$v_{tl-codon-i} = \mu \cdot c_{codon-i}. \tag{13}$$

Given these constraints, we minimize the total mass concentration of the modeled translation apparatus, consisting of ribosome, EF-Tu, EF-Ts, mRNA, GTP, GDP, and charged tRNAs (aa·tRNA),

$$\sum_{m \in C} MW_m \cdot c_m \quad , \tag{14}$$

where the molecule types together form the set $C$, $MW_m$ is the molecular weight, and $c_m$ is the concentration of molecule type $m$. We do not minimize the contributions of GTP and GDP, who participate in multiple other cellular processes [22] and whose concentrations are thus unlikely to be dominated by translation; their concentrations are consequently fixed to experimentally observed values in our simulations (see **Supplementary Note 3**).

Let $N$ be the set of reactions that together comprise translation. We also consider the dilution of the molecules involved in these reactions due to cellular volume growth at rate $\mu$:

$$\boldsymbol{S} \cdot \vec{v}(\vec{c}) - \mu \cdot \vec{c} = 0, \tag{15}$$

where $\boldsymbol{S}$ is the stoichiometric matrix for the reactions in $N$, $\vec{c}$ is a vector of the concentrations $c_m$, and $\vec{v}(\vec{c})$ is the corresponding vector of reaction rates $v_i$, with the concentration-dependent kinetics described above.

Thus, we solve the non-linear constrained optimization problem:

$$\min_{[\vec{c}]} \quad \sum_{m \in C} MW_m \cdot c_m \,. \qquad (16)$$

Subject to:

$$S \cdot \vec{v}(\vec{c}) - \mu \cdot \vec{c} = 0$$

$$v_{tl-codon-i} = \mu \cdot c_{codon-i} \quad \text{for } i\text{=1, …, 61}$$

$$v_{\text{term}} = \mu \cdot c_{\text{protein}}$$

We formulated the optimization problem in GAMS and used the BARON global solver[23] on NEOS Server[24] with 8 hours as the time limit to solve this problem.

### Supplementary Notes

#### 1. The coarse-grained optimization model by Klumpp *et al.*

Noting that the translation machinery includes not only ribosomes, but also other highly expressed proteins – most notably elongation factors[1] and tRNA synthetases – Klumpp *et al.*[20] argued that a full appreciation of the efficiency of protein synthesis requires the inclusion of the cost of these translation components. They suggested a phenomenological, coarse-grained model of cell growth, comprised of four proteome sectors, including a ribosomal (Rb) and a translation-associated (T) sector. Assuming co-regulation of the Rb and T sectors and fitting three phenomenological constants to the data, they were able to approximate the growth-rate dependence of ribosome concentration and elongation speed in *E. coli* [20]. However, the experimentally observed ratio between the protein concentrations in the T- and Rb-sectors, $\Phi_T/\Phi_{Rb}$ , deviates from the postulated constant ratio (see Fig. 3D in Ref. [20]), indicating shortcomings of this phenomenological theory.

Klumpp *et al.* also attempted to determine an optimal growth-rate dependence of the ratio between T- and Rb-sectors, by treating both proteome fractions as independent parameters when numerically optimizing the growth rate of their coarse-grained model cell. However, the results predicted a ratio $\Phi_T/\Phi_{Rb}$ that was substantially smaller than that observed (see Fig. 4C in Ref. [20]), indicating that translation in *E. coli* is either not organized optimally, or that the objective optimized by natural selection differs from the proteome allocation examined by Klumpp *et al.*. Comparing the objective functions used by Klumpp *et al.* and in the present work, we note that proteins make up 1/3 of the ribosome, but 2/3 of the ternary complex (by mass). Thus, the ternary complex appears much more expensive to the cell when considering protein mass than when considering total mass, explaining why optimization of protein allocation results in smaller predictions of the $\Phi_T/\Phi_{Rb}$ ratio[20,25].

#### 2. Ribosome states

The ribosome is the most central component of translation, and the ribosome states in our model are slightly different from those used in proteome partitioning models[4,11,20]. In this section, we will discuss the difference and the rationality of ribosome states in our model.

Briefly, our model contains active ribosomes and free ribosomes. **Active ribosomes** are bound to mRNA and actively involved in elongating peptide chains. **Free ribosomes** are responsible for translation initiation; they are available for binding to mRNA and comprise a subset of the inactive ribosomes in proteome partitioning models.

Proteome partitioning models distinguish between active ribosomes and inactive ribosomes. **Active ribosomes** have exactly the same meaning as in our model: they are engaged in elongation. At steady-state growth, the protein synthesis rate can be written as: $v_{\text{protein\_syn}} = \mu\, P = f_{\text{active}} \cdot k_{\text{eff}} \cdot R$, where $\mu$ is the growth rate, $P$ is the total protein concentration (measured in amino acids per volume), $f_{\text{active}}$ is the fraction of active ribosomes among total ribosomes, $k_{\text{eff}}$ is the turnover number of ribosomes during elongation, and $R$ is the concentration of ribosomes. By measuring $\mu$, $k_{\text{eff}}$, and the ratio between R and P (estimated through the RNA/Protein ratio and the fraction of rRNA in total RNA), Dai *et al.* estimated the fraction of active ribosomes as a function of growth rate[4]. In the view of protein partitioning models, the **inactive ribosomes** comprise all ribosomes not actively engaged in elongation. Inactive ribosomes include not only ribosomes available for initiation (**free ribosomes**), but also ribosomes that are unavailable for initiation (**unused, or deactivated,**

**ribosomes**) [4,11,20]. In this work, we modeled both initiation and elongation, and thus both free and active ribosomes (but not deactivated ribosomes) are included.

Our model is carefully built on first principles. All reactions are explicitly and exclusively constrained by reaction parameters and steady state growth; we avoid any empirical growth rate-dependent parameters, such as a growth rate-dependent fraction of active ribosomes or effective ribosome activity. In other words, our model contains only reactions for which we know why and how they occur. The mechanism leading to a fraction of deactivated ribosomes is not clear. Deactivated ribosomes facilitate faster transitions between growth environments that support different growth rates[26], a phenomenon that cannot be predicted with steady-state models such as ours. Moreover, the true fraction of deactivated ribosomes has not been measured experimentally. Thus, we did not attempt to predict the total concentration of ribosomes (including deactivated ribosomes), and only compared our predictions for active ribosome concentrations to experimental estimates.

#### 3. Impact of GTP and GDP concentrations on the predictions

GTP and GDP are involved in many intracellular reactions [22], and we thus do not expect to predict their concentrations in this translation model. In our model, GTP and GDP are involved in nucleotide exchange by elongation factor Tu [16] (see Methods). The concentrations of GTP and GDP may influence the rates of some reactions directly. In this section, we assess the impact of the assumed GTP and GDP concentrations on the predictions, examining three pairs of concentrations[5] resulting from growth of *E. coli* K-12 on different media. Note that GTP and GDP concentrations[5] and proteome data were collected for different strains of *E. coli* K-12 (NCM3722 and BW25113, respectively).

The GTP/GDP measurements were done for growth on acetate ($c_{GTP}$ = 1250 µM; $c_{GDP}$ = 18 µM), glycerol ($c_{GTP}$ = 2690 µM; $c_{GDP}$ = 23 µM), and glucose ($c_{GTP}$ = 4900 µM; $c_{GDP}$ = 680 µM); all three conditions also appear in our simulations. We first simulated growth on acetate and on glycerol with GTP and GDP concentrations measured for *E. coli* cells growing on the same media. Next, we replaced the GTP and GDP concentrations with the data for glucose and repeated the simulations (**Suppl. Fig. S1**). Despite the large differences in GTP and GDP concentrations, the results obtained are very similar. For both acetate and glycerol growth, geometric mean fold-errors (GMFE) are below 1.03 (Suppl. Fig. S1), *i.e.*, the predicted concentrations of the individual components of the translation machinery are on average less than 3% higher or lower in the two sets of predictions. Thus, GTP and GDP concentrations appear to have only a minor influence on the predictions. Because glucose is the reference condition for the protein expression data[1], we used the concentration of GTP and GDP for growth on glucose for all predictions in this study.

#### Supplementary Figures

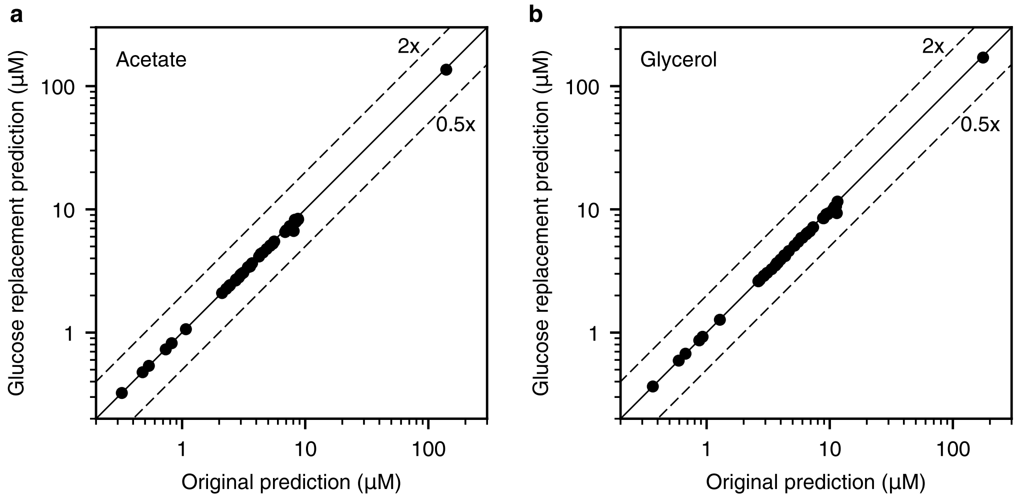


**Figure S1.** Impact of GTP and GDP concentrations on model predictions. **(a)** Growth on acetate (geometric mean fold-error *GMFE* = 1.026). **(b)** Growth on glycerol (*GMFE* = 1.029). Each datapoint represents the concentration of one model component (ribosome, EF-Tu, EF-TS, aa-tRNA). x-axes show predictions using the GTP and GDP concentrations measured for the corresponding medium; y-axes show predictions when instead assuming the GTP and GDP concentrations measured for growth on glucose. *GMFE* measures the mean deviation from the identity line on the log-log plot; *GMFE* = 1 indicates perfect identity. The very low *GMFE* values indicate that *in vivo* GTP and GDP concentration has a very small effect on our model.
